# Altered brain-wide auditory networks in *fmr1*-mutant larval zebrafish

**DOI:** 10.1101/722082

**Authors:** Lena Constantin, Rebecca E. Poulsen, Itia A. Favre-Bulle, Michael A. Taylor, Biao Sun, Geoffrey J. Goodhill, Gilles C. Vanwalleghem, Ethan K. Scott

## Abstract

Altered sensory processing is characteristic of several psychiatric conditions, including autism and fragile X syndrome (FXS). Here, we use whole-brain calcium imaging at cellular resolution to map sensory processing in wild type larval zebrafish and mutants for *fmr1*, which causes FXS in humans. Using functional analyses and graph theory, we describe increased transmission and reduced filtering of auditory information, resulting in network-wide hypersensitivity analogous to the auditory phenotypes seen in FXS.

---

Sensory processing phenotypes contribute to the diagnostic descriptors of psychiatric conditions such as FXS and autism spectrum disorder (ASD)^1^. However, the circuit-level causes of sensory phenotypes remain mysterious because of the technical challenges associated with observing entire functional networks at cellular resolution. Hence, there is a strong incentive to observe and model neural circuits *in vivo*, and to identify the network alterations that lead to sensory phenotypes in models of these disorders. To this end, we have performed whole-brain calcium imaging of visual and auditory processing in wild type (WT) zebrafish larvae, and larvae mutant for *fmr1*, which is responsible for FXS in humans^2^.

Using GCaMP6^3^ and light-sheet microscopy^4^, we performed volumetric imaging of both baseline and stimulus-evoked neuronal calcium activity at 2-4 Hz throughout the brains of 6-day-old zebrafish larvae **(Fig. S1a)**. We then segmented regions of interest (ROIs) generally corresponding to individual neurons and extracted the fluorescence traces from each ROI^5^ **(Fig. S1b)**. We first measured baseline activity in WT, heterozygous (het), and *fmr1*^−/−^ mutant larvae, and found similar numbers of calcium events across genotypes **(Fig. 1a**, **Table S1)**. To determine whether correlations among active neurons had increased, as occurs in the cortex of mid-adolescent *Fmr1*^−/−^ mice^6^, we calculated the correlation coefficients between all ROIs in each larva. Again, there were no significant differences in mean correlation **(Fig. 1b**, **Table S1)**. These results suggest that *fmr1*^−/−^ animals have roughly normal baseline neuronal activity and activity correlations at 6 days of age.

**Figure 1.**
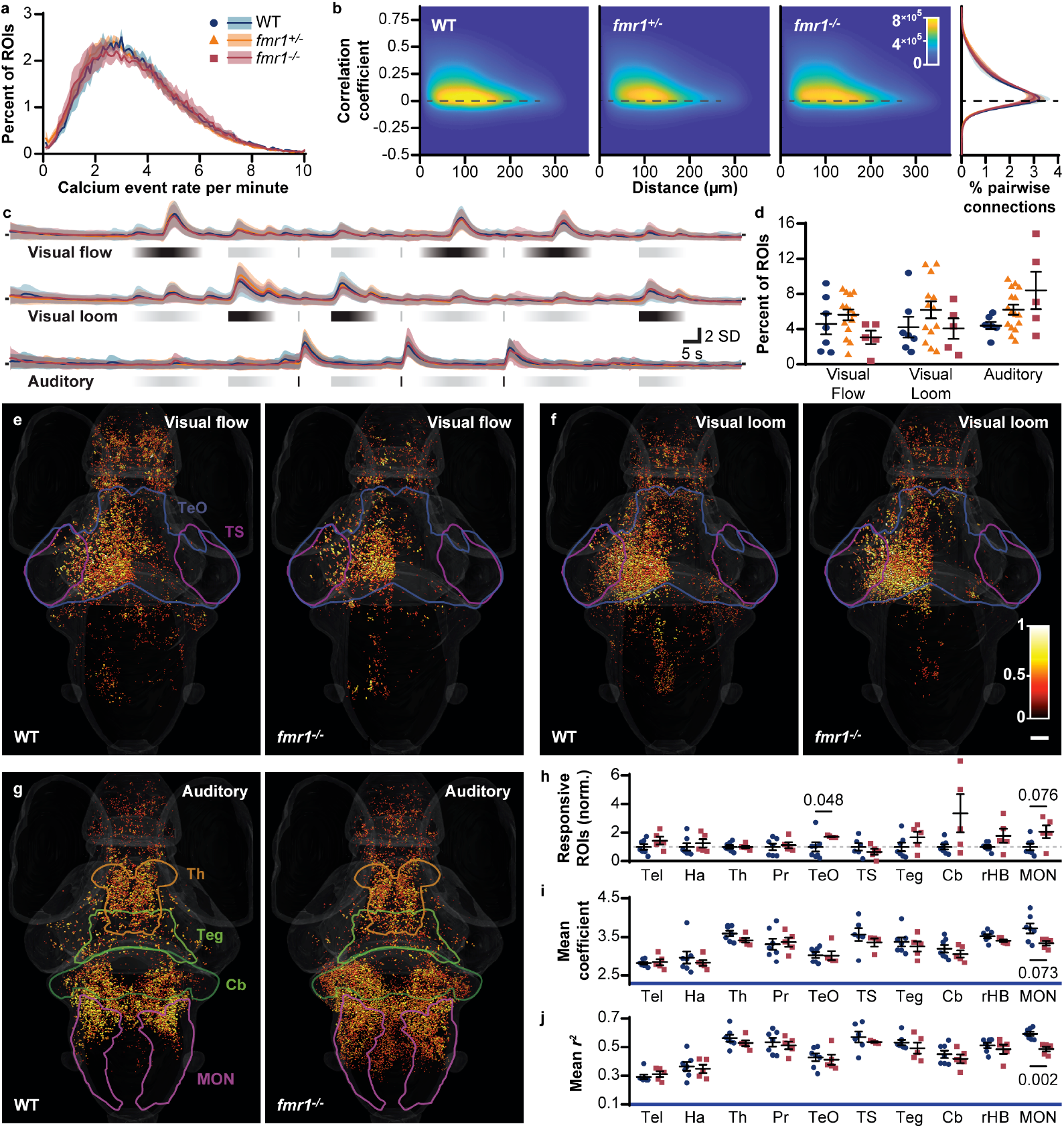
Brain-wide baseline and sensory-evoked neuronal activity. Distribution of brain-wide calcium event rates **(a)** in WT, *fmr1*^+/−^, and *fmr1*^−/−^ larvae at baseline (mean ± s.e.m.). Heat map **(b)** of the regression coefficient between pairs of ROIs as a function of Euclidean distance between the ROIs, with a distribution of all coefficients (right) showing no marked differences among the genotypes. Z-scored activity traces **(c)** of all ROIs responding to stimuli in each genotype (mean ± SD shown for all ROIs with a regression coefficient > 2 SD and > 0.1 *r*^*2*^ value to the relevant stimulus). Percent of all ROIs **(d)** across the brain responding to each stimulus (mean ± s.e.m.; each data point represents one animal, n=7 WT and 5 *fmr1*^−/−^). **(e-g)** Brain-wide responses to each stimulus across all animals (28.6% of WT responses are omitted, at random, to account for the use of 7 WT versus 5 *fmr1*^−/−^ animals). Spot colour represents response strength (correlation coefficient) and spot diameter depicts response fidelity (*r*^*2*^ value). Scale bar **(f)**, is 50 microns and applies to panels **e-g**. Ratio of ROIs **(h)** in WT versus *fmr1*^−/−^ responding to the auditory stimulus across various brain regions (mean ± s.e.m., one data point per animal). Mean regression coefficients **(i)** and *r*^*2*^ values **(j)** of ROIs responding to auditory stimuli (regression coefficient > 2 SD and *r*^*2*^ > 0.1) across various brain regions in WT and *fmr1*^−/−^ larvae (mean ± s.e.m.). Brain regions showing trends of interest are outlined **(e-f)**. Cb, cerebellum; Ha, habenulae; MON, medial octavolateralis nucleus; Pr, pretectum; rHB, remaining hindbrain with the Cb and MON excluded; Teg, tegmentum; Tel, telencephalon; TeO, optic tectum; Th, thalamus; TS, torus semicircularis. *P*-values (t-test with Mann-Whitney correction) are shown for all cases where p < 0.1.

Humans with FXS show deficits in visual motion detection^7–9^ and hypersensitivity to auditory stimuli^10,11^. To judge whether *fmr1*^−/−^ larvae have similar sensory phenotypes, we presented two visual stimuli (moving lines that provide visual flow and a looming disk) and one auditory stimulus (white noise at 84 decibels (dB)) to larvae while performing calcium imaging. To identify responsive ROIs, we used multivariate linear regressions to compare each z-scored calcium trace to regressors for the three stimuli. For each ROI, the regression coefficient provided a readout of response strength, while the *r*^*2*^ value represented the fidelity of its response to the stimulus. We calculated the mean z-scored fluorescent trace of responsive ROIs for each genotype to estimate the number and quality of responses across the whole brain **(Fig. 1c)** and the proportion of responsive ROIs per larva **(Fig. 1d)**. These analyses showed similar fluorescent activity profiles for all stimuli and similar proportions of visually responsive ROIs, but a trend towards more auditory responsive ROIs in *fmr1*^−/−^ larvae.

To explore whether the *fmr1* mutation altered brain responses in specific brain regions, we mapped each responsive ROI back to its anatomical position in a reference brain^12^. This allowed for the quantification of ROI number, response strength, and response fidelity for different brain regions. Brain-wide, the distributions and strengths of the responses were similar for visual flow **(Fig. 1e**; **Table S2)** and visual loom **(Fig. 1f**; **Table S2)**, and these parameters were quantitatively similar across individual visually responsive brain regions **(Fig. S2**; **Table S2)**. For the auditory stimulus, responses were distributed more broadly in *fmr1*^−/−^ larvae **(Fig. 1g)**, especially in the cerebellum, hindbrain, and medial octavolateral nucleus (MON, homologous to the cochlear nucleus). There was also a trend towards more auditory responsive ROIs in several brain regions **(Fig. 1h)**. In most regions, the average strengths **(Fig. 1i**; **Table S2)** and fidelities **(Fig. 1j**; **Table S2)** of responses were similar, but ROIs in the MON of *fmr1*^−/−^ larvae were, although more numerous, weaker and noisier. In summary, our initial analyses provided no compelling evidence for network-level alterations in visual motion processing in *fmr1*^−/−^ larvae. Auditory processing appeared to be more abundant and dispersed in *fmr1*^−/−^ animals, but these results were statistically marginal, and the use of a single auditory stimulus limited our ability to draw functional conclusions about alterations in the network.

To delve deeper into the auditory phenotype, we designed an auditory-only stimulus train comprising an ascending amplitude ramp, played twice, and twelve volumes of a brief auditory stimulus, played three times each. The overall distribution of responsive neurons **(Fig. 2a)** was similar to that observed with our simple auditory stimulus **(Fig. 1g)**, with a larger number of more broadly distributed auditory ROIs. The mean z-scored fluorescent traces of brain-wide auditory ROIs were similar in WT and *fmr1*^−/−^ larvae, but there was a tendency toward greater responsiveness in *fmr1*^−/−^ larvae to weak stimuli **(Fig. 2b)**.

**Figure 2.**
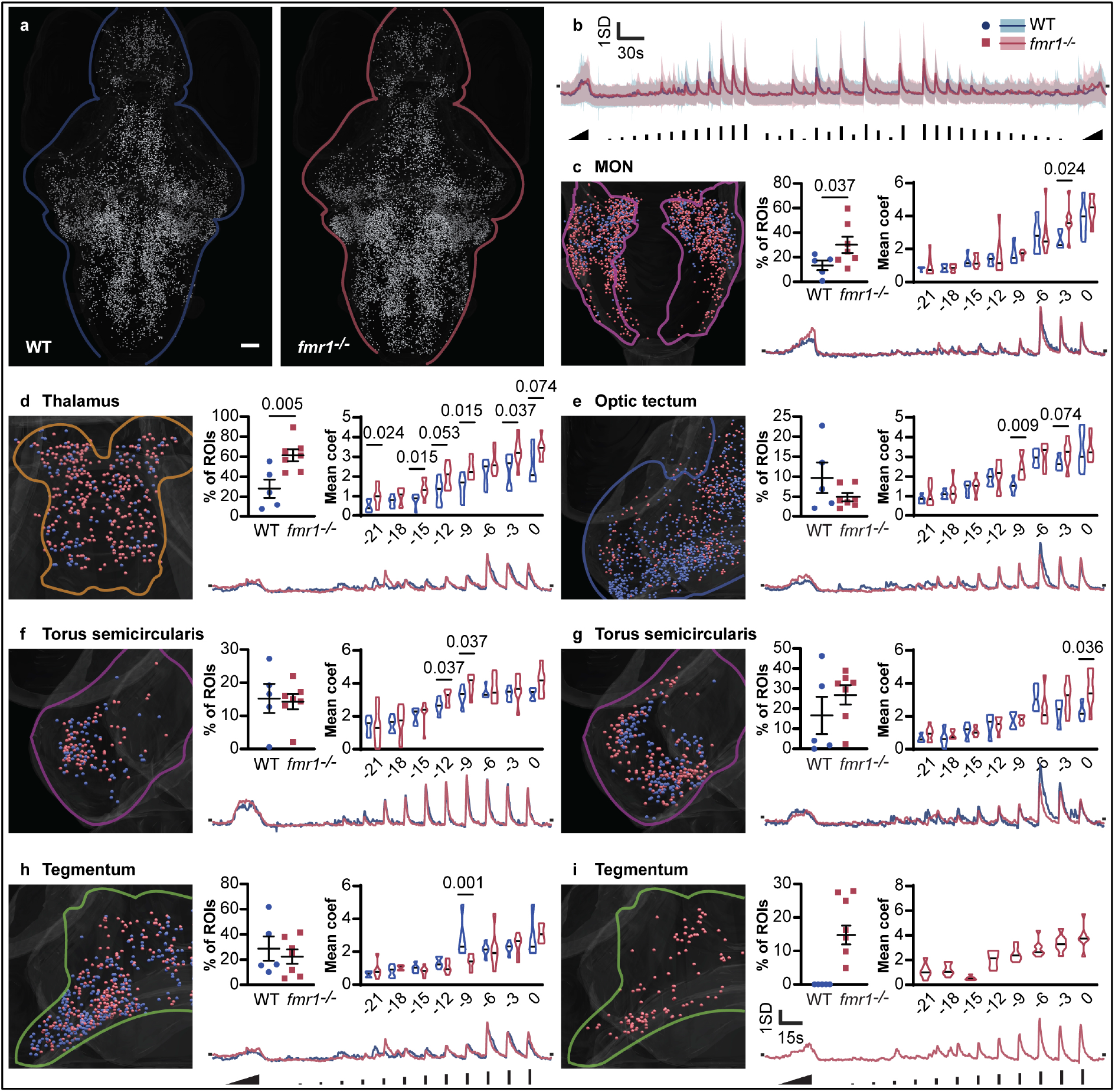
Regional responses to a complex auditory stimulus train. **(a)** Auditory responsive ROIs in WT (n=5) and *fmr1*^−/−^ larvae (n= 7, 28.6% of ROIs omitted throughout this figure to provide equivalent images to WT). Mean z-scored activity trace **(b)** of all responsive ROIs (with stimulus timing and volume represented at the bottom). For each of seven functional clusters across five brain regions **(c-i)**, the distribution of responsive ROIs (left), percentage of all ROIs belonging to the cluster and mean response strength to each of eight stimulus volumes (top right), and mean z-scored activity trace during the first amplitude ramp and first twelve discrete volumes (bottom right). Volumes are represented in decibels (dB) below full volume, and represent the 5^th^-12^th^ volumes in the entire stimulus train.

We next performed an in-depth analysis of auditory responses across numerous brain regions (see **Table S3)** previously shown to have a role in sensory processing in zebrafish larvae^5^. First, we applied k-means clustering on the time series, with the city block metric, to identify functionally distinct categories (clusters) of ROIs with consistent and characteristic responses to our auditory stimuli. For each brain region and cluster, we measured the proportion of all ROIs in each brain region that belonged to the relevant cluster, compared their mean z-scored fluorescent traces, and quantified their average response strengths at each auditory volume **(Table S3)**. Five brain regions showed trends or significant or differences in *fmr1*^−/−^ versus WT **(Fig. 2c-i)**, and each cluster in each region and genotype was represented by at least 80% of the larvae, showing that the observed effects were not products of artifacts in individual animals. Starting with the MON, the first brain region to receive auditory input from the vestibulocochlear nerve (cranial nerve VIII)^13^, we identified a single functional cluster with significantly more ROIs in *fmr1*^−/−^ larvae. This difference was the result, at least in part, of auditory responses extending more caudally into the MON (**Fig. 1g**; **Fig. 2a, c)**. In the thalamus (**Fig. 2d)**, there were significantly more auditory responsive neurons in *fmr1*^−/−^ larvae, combined with an increase in the response strength of these ROIs across a range of volumes. The tecta of *fmr1*^−/−^ animals, in contrast, had fewer auditory responsive neurons, and showed similar response profiles across the genotypes **(Fig. 2e)**. Two separate functional clusters emerged in the torus semicircularis. The first cluster **(Fig. 2f)** was incrementally sensitive to a wide range of volumes, and showed no significant differences in the proportions, response strengths, or response traces between genotypes. The second, less sensitive cluster of the torus semicircularis **(Fig. 2g)** responded to stronger volumes. In *fmr1*^−/−^ animals, these ROIs were more numerous and had significantly elevated response strengths at higher volumes. A cluster with similar response characteristics was present in the tegmentum **(Fig. 2h)**, where there were no pronounced differences between genotypes. A second tegmental cluster was identified exclusively in *fmr1*^−/−^ animals **(Fig. 2i)**, with response strengths that better reflected the stimulus intensity across a range of volumes.

We next performed graph theory to gauge the impacts of the *fmr1* mutation on the overall brain-wide auditory network. For the WT and *fmr1*^−/−^ datasets, we generated sets of 132 and 134 nodes, respectively, that represented the functional units of audition across the ten brain regions of interest **(see Online Methods)**. The flow of information among nodes was then modelled by calculating the correlation coefficient across all pairs. Matrices representing correlations among all pairs of nodes **(Fig. 3a)** showed stronger correlations in *fmr1*^−/−^ larvae across all volumes tested (the 6^th^ to 10^th^ out of 12 ascending volume stimuli are shown), demonstrating higher overall network correlation. In *fmr1*^−/−^ larvae, we found enhanced density, which measures the portion of each node’s possible edges that are strongly correlated, across the network as a whole, a result that was consistent across all volumes and correlation thresholds **(Fig. 3b)**. Participation, a measure of each brain region’s correlation with other brain regions’ nodes, was increased in *fmr1*^−/−^ larvae for all regions tested except for the torus semicircularis **(Fig. 3c)**. We next mapped all the nodes spatially based on the centres of gravity for the ROIs composing them and represented strong pairwise correlations as edges **(Fig. 3d**, showing the 5^th^ to 12^th^ volumes). Combined, these data suggest stronger brain-wide correlations in the brains of *fmr1*^−/−^ larvae versus WT, and greater transmission of information in the early ascending auditory pathway to other brain regions, especially at lower volumes.

**Figure 3.**
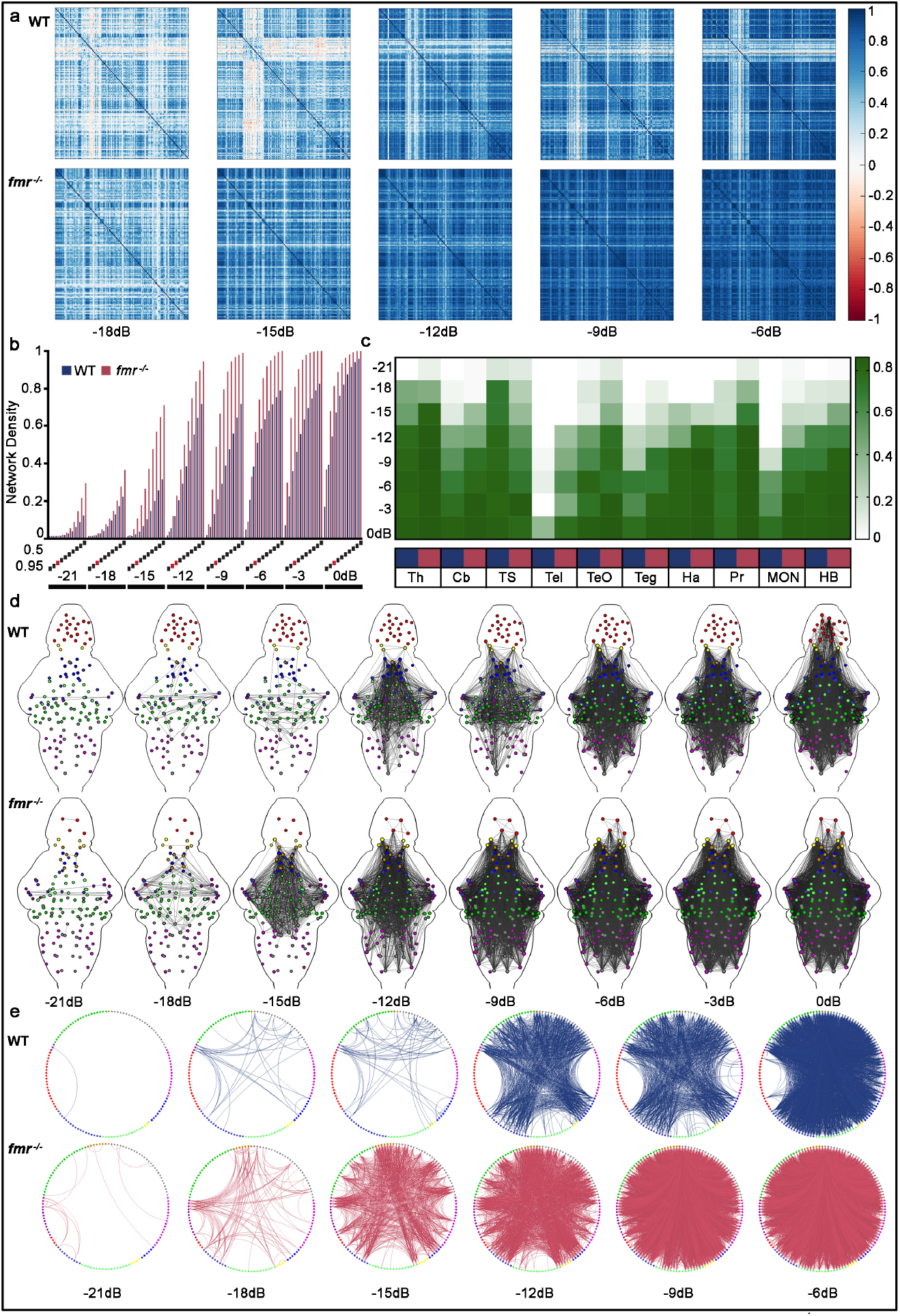
Functional brain-wide auditory networks in WT and *fmr1*^−/−^ larvae. Correlation matrices **(a)** showing pairwise correlation strength (colour map, right) across all pairs of nodes in WT (top) and *fmr1*^−/−^ (bottom) larvae. Volumes are annotated as decibels from full volume (dB). Network density **(b)** as a function of correlation coefficient thresholds (0.95 to 0.5 in 0.05 increments) and volume (the 5^th^-12^th^ volumes in our stimulus train). The 0.85 correlation coefficient threshold (red dash) was selected for subsequent analyses. **(c)** The mean participation coefficient (proportion of nodes with edges to other brain regions) for each region (x-axis) across nine volumes (y-axis), with the colour scale indicated on the right. Blue is WT and red is *fmr1*^−/−^. **(d)** Graphs of the brain-wide auditory network, showing edges exceeding a correlation coefficient of 0.85. Node colour indicates brain region: Cerebellum, dark green; habenulae, yellow; MON, magenta; pretectum, light blue; hindbrain without the cerebellum and MON, grey; tegmentum, light green; optic tectum, blue; telencephalon, red; thalamus, orange; torus semicircularis, dark magenta. Circle plots **(e)** of strongly correlated edges (above 0.85 correlation coefficient) between nodes (circles) located in different brain regions (coloured as above) for different auditory volumes in WT and *fmr1*^−/−^.

To clarify which regions were correlated more strongly in *fmr1*^−/−^ animals, we organised nodes by brain region and represented the strongly correlated edges at different stimulus volumes **(Fig. 3e)**. This analysis revealed more functionally correlated regions at lower volumes in *fmr1*^−/−^ animals, with a greater number of brain regions engaging earlier. In WT larvae, the first edges formed between the torus semicircularis at −21 dB, with subsequent increases in volume leading to interactions between the torus semicircularis and the tectum, hindbrain, and pretectum, in addition to edges between the thalamus and pretectum. As the volume increased further, the cerebellum engaged, and by −9 dB, all brain regions of interest were engaged. In contrast, the *fmr1*^−/−^ larvae showed strong edges between the torus semicircularis and the tectum at −21 dB, with nearly all brain regions by −18 dB.

Combined, alterations in the distributions and response profiles of neurons within brain regions **(Fig. 2)**, and in the correlations between the neurons’ representative nodes across the brain **(Fig. 3)**, have revealed fundamental network properties of an *fmr1* auditory phenotype. During auditory processing, information from the vestibulocochlear nerve is first received by the MON (homologous to cochlear nuclei of mammals) and mediodorsal hindbrain (presumed to develop into the secondary octaval population in mammals^13^), which project to the torus semicircularis (homologous to the inferior colliculus) *en route* to the thalamus^13,14^. Finally, the dorsomedial telencephalon, which is homologous to the mammalian amygdala, and the thalamus are reciprocally connected^14^.The torus semicircularis also relays auditory information to the deeper layers of the optic tecta (homologous to the superior colliculus^11^) and tegmental nuclei, which form a part of a descending inhibitory reflex circuit with the hindbrain and spine^15^.

Our brain-wide analysis at cellular-resolution shows some of the population-scale properties of the *fmr1* phenotype. Specifically, we find greater responsiveness to auditory stimuli (manifested as a greater number of responsive neurons) in *fmr1*^−/−^ larvae early in this pathway (in the MON), although the response profiles of these neurons are not markedly different from those in their WT counterparts **(Fig. 2c)**. Downstream of the MON, the torus semicircularis shows more responsive neurons, especially for the functional cluster with responses like those in the MON **(Fig. 2g)**. Later in the pathway, we see greater numbers of neurons with stronger auditory responses (in the thalamus, **Fig. 3d**) and an entire functional cluster with MON-like responses that exist only in *fmr1*^−/−^ larvae (in the tegmentum, **Fig. 3i**). These data point towards the greater transmission of information, with less modulation and refinement, in the auditory pathways of *fmr1*^−/−^ larvae.

At the network level, our graph analysis represents brain-wide information flow based on data from individual neurons. This reveals higher correlations among virtually all auditory responsive brain regions **(Fig. 3a)**, and notably between regions early (MON) and late (thalamus) within the pathway **(Fig 3b, e)**. Auditory hypersensitivity is a characteristic of people with FXS^16^, and this is reflected by a roughly 6dB (or 4-fold) leftward shift in the network responsiveness of *fmr1*^−/−^ larvae, regardless of the metrics used **(Fig. 3 a-c, e)**. Furthermore, hyperconnectivity of the thalamus, based on functional MRI studies, is consistently reported in ASD^17–19^, which shares high comorbidity with FXS^20^. We show that functional hyperconnectivity in the thalamus likely arises from the combined effects of more plentiful and stronger neuronal responses **(Fig. 2d)** and stronger links to other brain regions (orange nodes, **Fig. 3e**). Beyond the thalamus, hypersensitivity was observed in the form of more numerous responses to auditory stimuli in the tegmentum, and engagement of the tegmentum with the rest of the auditory network at lower volumes.

In summary, we report an analysis of auditory processing in *fmr1*^−/−^ zebrafish larvae that spans individual neurons, local populations, and brain-wide networks. Across these scales, our results indicate that auditory information is transmitted more strongly, and with less modification and refinement, as it passes through the auditory processing pathway in *fmr1*^−/−^ animals. This results in greater and less discriminant activity later in the pathway and in stronger correlations across the network at lower stimulus volumes. This reveals an overarching mechanism for auditory hypersensitivity in these animals, with implications for humans with FXS, and provides departure points for targeted functional studies of the underlying circuit-level mechanisms.

## Acknowledgements

We thank Emmanuel Marquez-Legorreta for his intellectual expertise on the looming stimulus, and the University of Queensland’s Biological Resources aquatics team for animal care. Support was provided by the Simons Foundation Autism Research Initiative through Pilot (399432) and Explorer (336331) awards, an NHMRC Project Grant (APP1066887), two ARC Discovery Project Grants (DP140102036 & DP110103612), and an ARC Future Fellowship to E.K.S.; an EMBO Long-Term Fellowship to G.C.V.; a fellowship from the Human Frontier Science Program (LT000146/2016) to M.A.T.; and an Australian Postgraduate Award to R.E.P.

## Author Contributions

Conceptualization, E.K.S.; Methodology, L.C., R.E.P., G.C.V., I.A.F-B and M.A.T.; Investigation, L.C., R.E.P., G.C.V.; Animal Colony Maintenance, L.C. and B.S.; Formal Analysis, L. C., R.E.P., G.C.V.; Data Curation, L.C. and R.E.P.; Writing–Original Draft, L.C. and E.K.S.; Writing–Review and Editing, R.E.P. and G.C.V.; Figure Construction, R.E.P. and L.C.; Funding Acquisition, E.K.S.; Resources, E.K.S. and G.J.G.; Supervision, E.K.S. and L.C.

## Methods

### Animal models

All work was performed in accordance with the University of Queensland Animal Welfare Unit (approval SBMS/378/16). Adult zebrafish (*Danio rerio*) were reared and maintained in a Tecniplast zebrafish housing system under standard conditions using the rotifer polyculture method for early feeding 5 to 9 days post fertilization (dpf). Embryos were reared in embryo medium (1.37 mM NaCl, 53.65 μM KCl, 2.54 μM Na_2_HPO_4_, 4.41 μM KH_2_PO_4_, 0.13 mM CaCl_2_, 0.16 mM MgSO_4_, and 0.43 mM NaHCO_3_ at pH ~7.2) at 28.5 °C on a 14-hour light: 10-hour dark cycle. To generate the experimental cohort, zebrafish carrying the *fmr1*^*hu2787*^ allele^1^ were bred to zebrafish carrying the *elavl3:H2B*-*GCaMP6* transgene^2^. The fast variant of GCaMP6, *elavl3:H2B*-*GCaMP6f*, was used to capture baseline and auditory sensitivity activity. The slow variant, *elavl3:H2B*-*GCaMP6s* was used for multisensory-evoked experiments. Zebrafish mutant transgenic animals were out-crossed for four generations to the *Tupfel long fin nacre* wild-type strain. Genotyping for *fmr1* was performed as previously described^3^. We generated the experimental cohort by inter-crossing the fourth generation of zebrafish heterozygous for both *fmr1*^*hu2787*^ and *elavl3:H2B*-*GCaMP6* to produce clutches with a Mendelian ratio of 1:2:1 (wild-type (WT):*fmr1*^+/−^: *fmr1*^−/−^). Larvae were pre-screened for GCaMP6 expression under a fluorescence microscope at 3dpf. Experiments and quality control were performed blinded to genotype.

### Animal preparation for calcium imaging

Larvae at 6 dpf were embedded upright in 1.5% low melting temperature agarose (Sigma, A9045) inside of a custom-built imaging chamber. Imaging chambers were composed of a 3D-printed base (24 × 24 mm) with four posts (3 × 3 × 20 mm) raised along the four corners of the platform. The four outward faces of the chambers were fixed with a glass coverslip (20 × 20 mm, 0.13-0.16 mm thick). Individual larvae were mounted onto a raised platform (11 × 11 × 6 mm) within each chamber. The platform was no closer than 3 mm from any of the glass surfaces. Chambers were filled with embryo medium once the surrounding agarose had set.

### Experimental setup for calcium imaging

Visual stimuli were displayed on a 75 × 55 mm LCD generic PnP monitor (1024 × 768 pixels, 85 Hz, 32-bit true colour) positioned 35 mm lateral to the larva. The monitor was covered by a coloured-glass filter (Newport, 65CGA-550) with a cut-on wavelength of 550nm. Auditory stimuli were played from a miniature speaker (Dayton Audio DAEX-9-4SM Skinny Mini Exciter Audio and Haptic Item Number 295-256) fixed to the glass surface of the imaging chamber posterior to the larva. The miniature speaker was wired to a Dayton Audio DA30 2 x 15W Class D Bridgeable miniature amplifier.

### Stimulus trains for calcium imaging

For multisensory experiments, we presented three sensory stimuli to each larva three times in a semi-randomised order. The stimuli had a minimum interstimulus interval of 3 seconds. Following 15 seconds of acclimatisation to laser scanning, the brain was imaged for 40 seconds at rest and then for 3 minutes and 12 seconds over the course of the multisensory stimulus train. We presented visual stimuli at a resolution of 1024 × 768 pixels at 20 frames per second. The two visual stimuli were moving vertical bars (visual flow) and an expanding disk (visual loom). The visual flow stimulus comprised eight bars, 128 pixels in width, moving in a caudal to rostral direction at a speed of 21.3° per second for 4 seconds. Prior to each visual flow, a 12 second linear dimming from white to black-and-white bars occurred, and similarly, a black-and-white to white 12 second linear brightening occurred after each visual flow (total stimulus duration 28 seconds). The visual loom consisted of a black 4-pixel diameter circular disk that exponentially expanded (Simple, Fastest; easing of −80 designed on Adobe Animate v18.0) to an 812-pixel diameter circular disk over 6 seconds. A linear brightening from black to white over a 12 second duration followed each visual loom (total stimulus duration 18 seconds). The auditory stimulus was created using the professional audio software, Live! (Ableton) and was normalised to 0 decibels (dB) relative to full scale. It comprised 1 second of white noise with a 2 millisecond rise/fall time. The sound level of white noise at full scale was 84 dB (background noise 40-45 dB) and was measured using a digital sound level meter 0.5-inch electret condenser microphone (ST-85A) positioned just above the imaging platform (where the larvae would be). The total length of the multisensory-evoked stimulus train was 4 minutes.

For auditory sensitivity experiments, we presented two 30-second white noise amplitude ramps (to 0 dB), and twelve discrete volumes of white noise at 3 dB intervals. We played each discrete volume once in three blocks, with an inter-stimulus interval of 14 seconds within blocks and an inter-block interval of 30 seconds. Following 15 seconds of acclimatisation to laser scanning, the brain was imaged for 90 seconds during which the first amplitude ramp was presented. The first block comprised the twelve volumes with increasing amplitudes (−33, −30, −27, −24, −21, −18, −15, −12, −9, −6, −3, 0 dB). The second block was quasi-randomised (−21, −27, −12, −33, −9, −18, −6, −24, 0, −15, −30, −3 dB) and the third block had decreasing amplitudes (0, −3, −6, −9, −12, −16, −20, −24, −27, −30, −33 dB). The second amplitude ramp was presented at the end of the stimulus train. The total length of the auditory stimulus train was 11 minutes and 45 seconds.

### Calcium imaging

In vivo GCaMP6 imaging was performed on a custom-built light-sheet microscope. The microscope has been previously described^4^, including a line diffuser used to reduce striping artifacts^5^. For multisensory experiments, to avoid the laser paths streaming directly into the eyes (which would interfere with the perception of visual stimuli), we blocked the side laser plane and restricted the front laser plane to the area between the eyes using custom-made sheets. For baseline and auditory sensitivity experiments, we used both the side and front laser planes. The exposure time for all experiments was 10 milliseconds. The captured images were binned 4x, yielding a final image resolution was 640 × 540 pixels at 16-bit in a tagged image file format. For baseline and auditory sensitivity experiments, 25 transverse sections at 10 μm increments were recorded at four brain volumes per second for 10 minutes. Similar numbers of ROIs were detected per larvae (WT = 16,593 ± 1,165 across 7 animals *fmr1*^+/−^ = 17,077 ± 1,316 across 10 animals; *fmr1*^−/−^ = 17,321 ± 670 across 5 animals (mean ± s.e.m.)). For multisensory experiments, 50 transverse sections at 5 μm increments were sampled at two brain volumes per second for 4 minutes and 7 seconds. Again, similar numbers of ROIs were detected per larvae (WT = 45,015 ± 4,177 across 5 animals; *fmr1*^+/−^ = 49,768 ± 2,141 across 17 animals; *fmr1*^−/−^ = 40,123 ± 3,137 across 7 (mean ± s.e.m.)).

### Analysis of calcium activity

We analysed larvae that met the following four criteria: (1) showed robust responses to the first visual loom and auditory stimuli (6 of 77 excluded); (2) survived to the end of the imaging session (1 of 77 excluded); (3) the number of regions of interest (ROIs) detected within any of the brain regions of interest were within an order of magnitude of the median (11 of 77 excluded); (4) contained WT or *fmr1*^−/−^ larvae (that is, not *fmr1*^+/−^ exclusively) during the imaging session (imaging sessions were between 4 to 12 minutes; 7 of 77 excluded). We used the same animals for the baseline and auditory sensitivity experiments (*n* = 23), and a separate animal cohort for the multisensory experiments (*n* = 29). We measured baseline activity first, followed by auditory sensitivity.

For larvae passing the inclusion criteria, we separated their four-dimensional imaging stacks (time, x, y, z) into individual time series for every x-y plane using ImageJ v1.52c. Each plane was then motion corrected using the Non-Rigid Motion Correction (NoRMCorre) algorithm^6^. ROIs, and their corresponding calcium traces, were extracted, de-mixed and denoised using the CaImAn package^6^ as previously described^4^. Calcium traces of all animals and all planes were subsequently pooled per genotype and z-transformed for further analysis using MATLAB v9.5. For baseline activity, we computed the correlation between all pairs of ROIs and used their spatial localization to calculate Euclidean distances. We estimated calcium event rates by detecting the peaks in each trace that were separated by at least two seconds and that increased locally by at least 1 SD above the baseline. For evoked activity, we modelled calcium traces using multivariate linear regression against regressors (using a typical GCaMP6 response) that corresponded to the timing of the relevant stimulus types. For multisensory-evoked experiments, we built three regressors for each of the three presentations of visual flow, visual loom, and auditory stimuli. Likewise, for the auditory sensitivity experiments, we built twelve regressors for each of the repetitions of volume.

### Thresholding and calculating responsiveness

For the multisensory dataset, we defined an ROI as responsive to a particular stimulus if it had a regression coefficient 2 SD above the mean of all regression coefficients and had an *r*^*2*^ value greater than 0.1 (26^th^ percentile) in the WT group. WT thresholds were applied to *fmr1*^+/−^ and *fmr1*^−/−^ groups. Specifically, the regression coefficient thresholds 2 SD above the mean for visual flow, visual loom, and auditory were respectively 2.0462, 2.0920, and 2.2894.

For the auditory sensitivity dataset, we defined an ROI as responsive to a particular stimulus if it had a regression coefficient greater than 0 and had an *r*^*2*^ value greater than 0.05 (80^th^ percentile). Lower thresholds were used for the auditory sensitivity dataset (compared to the multisensory dataset) because we wanted to detect all responses to all auditory stimuli, including low volumes, and to accommodate a longer stimulus train. Given the complexity of the auditory stimulus train, and the low stringency of the thresholds, non-auditory signals had been pooled with the signal. To improve the signal-to-noise ratio without increasing the stringency of the thresholds (considering that we wanted all auditory categories including weak responses), we excluded timepoints in individual fish that startled at a particular volume, as these represent motor responses rather than auditory responses. In the WT group, we removed 50 frames flanking the first −6 dB volume in fish 1, the first −16 dB volume in fish 2, the second 0 dB volume in fish 3, and the third −27 dB volume in fish 5. In the *fmr1*^−/−^ group, we removed 50 frames flanking the first −6 dB volume in fish 4 and the second −33 dB volume in fish 7. We subsequently applied k-means clustering on the time series from the ten brain regions of interest to produce 5 components per genotype. All non-auditory clusters were excluded, and the remaining auditory clusters of all animals and all planes were pooled together for each region per genotype. Two different response profiles emerged in the tegmentum and torus semicircularis that remained in two distinct clusters. As k-means forces all ROIs to belong to a cluster, we removed ROIs with a low correlation to the mean of each cluster (correlation > 0.2) to remove additional noise. The resulting ROIs were classified as auditory responsive in the auditory sensitivity dataset.

To calculate the percent of ROIs responsive to a particular stimulus, ROI numbers above the regression coefficient and *r*^*2*^ thresholds were normalised to the total number of ROIs detected in the whole brain or relevant brain regions. To detect changes in the distribution of regression coefficients, the mean regression coefficients were calculated for ROIs with an *r*^*2*^ above threshold (0.1 or 0.05 depending on the dataset). Similarly, to detect changes in the distribution of *r*^*2*^ values, the mean *r*^*2*^ was calculated for ROIs per larvae with a regression coefficient above threshold (+ 2 SD of the WT or 0 depending on the dataset).

### Registration and visualisation of calcium activity

Once the motion corrected individual x-y planes were time averaged, we used the three-dimensional stacks of all fish included in a particular dataset to build a common template with Advanced Normalization Tools^7^. We then registered the template to the *Elavl3-H2BRFP* line on the Z-brain atlas^7^. The resulting warps were applied to the centroids of all ROIs for each larva, which were then placed into the 294 brain regions defined by the Z-brain atlas. Based on previous work^4^, we selected ten brain regions to investigate in greater detail. These were the cerebellum, habenula, medial octavolateralis nucleus, pretectum, remaining hindbrain (without the cerebellum and medial octavolateralis nucleus), optic tectum, tegmentum, telencephalon, thalamus, and torus semicircularis. We visualised the spatial information and classified activity of each ROI using the Unity Editor (https://unity3d.com/). Specifically, for the multisensory dataset, we represented ROIs as spheres with their diameter representing their *r*^*2*^ value (1 + coefficient of determination (*r*^*2*^) × 5 in μm), and colour based on their regression coefficient value. For the auditory sensitivity dataset, spot size and colour were uniform. The template brain, upon which we overlaid this information, was generated by creating an isosurface mesh over the combined masks of the diencephalon, mesencephalon, rhombencephalon, eyes, and telencephalon (from the Z Brain Atlas) using ImageVis3D.

### Graph theory

We defined the number of nodes by performing k-means clustering on the 3-dimensonal coordinates of all ROIs within each of the ten brain region per genotype into *k* number of clusters. *k* was chosen as the largest number (starting at 20 + number of ROIs/1000) at which at least 10 ROIs from at least 3 different fish were retained. This produced a total of 132 nodes in the WT and 134 in the *fmr1*^−/−^ cohort that were represented by at least 3 fish per genotype. We generated the mean z-scored fluorescent response from all the ROIs belonging to these nodes across all fish. Those mean responses were used to build correlation matrices for each genotype and binarized with a threshold of 0.85 correlation. The binary matrices were used to build undirected graphs. The Brain Connectivity Toolbox^8^ was used to calculate network measurements of the graph from each genotype, such as the density or the participation coefficient between brain regions. Circle plots were produced using the circularGraph toolbox (https://au.mathworks.com/matlabcentral/fileexchange/48576-circulargraph).

### Statistical analyses

Significance was tested using Mann-Whitney tests to compare two groups, and unpaired Kruskal-Wallis tests with Dunn’s multiple comparisons to compare three groups, in GraphPad Prism v8.0.1. Two-tailed tests were performed on the multisensory-evoked and baseline datasets. One-tailed tests (anticipating the trends of the multisensory-evoked dataset) were performed on the auditory sensitivity dataset to confirm that the same trends and directions observed in the multisensory-evoked were present in the auditory sensitivity dataset.

### Data and software availability

Data and software will be made available upon request.

## Supplementary Figures and Tables

**Supplementary figure 1.**
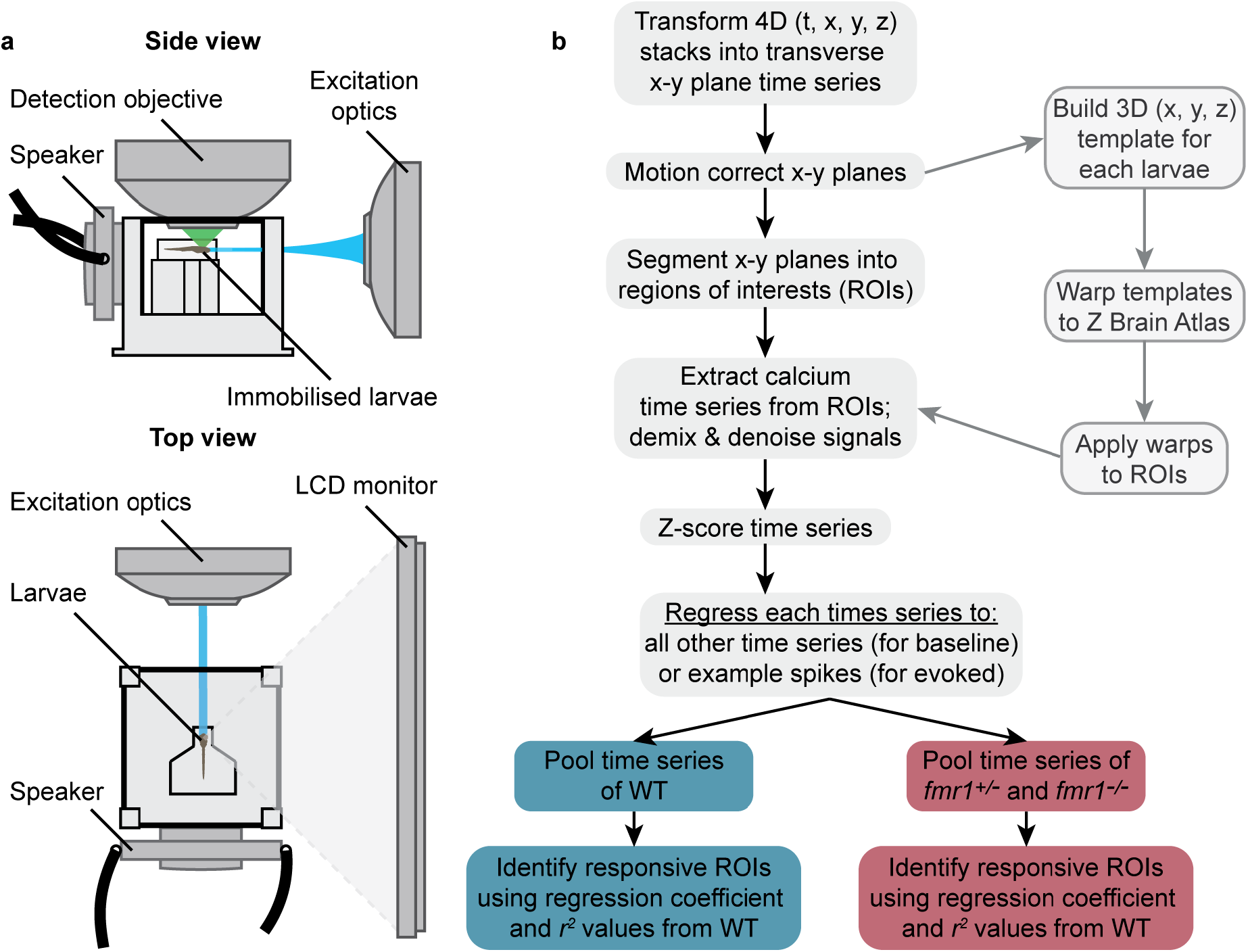
Imaging setup and neuroinformatics workflow. Schematic **(a)** of imaging setup used to collect baseline and evoked brain activity from 6 dpf zebrafish larvae. All larvae expressed the calcium indicator GCaMP6. For baseline activity, no stimuli were presented, although light from the excitation optics was present. For evoked activity, two visual and one auditory stimulus were presented three times in a quasi-random order. Visual stimuli were displayed perpendicular to the right eye of the larvae on an LCD monitor. Sound was played using a rear-mounted miniature speaker. Concurrently, calcium activity was recorded at 2-4 volumes per second. Summary of the neuroinformatics workflow **(b)** used to extract information related to calcium activity (left arm) and location (right arm) from four-dimensional (4D) imaging stacks. Briefly, 4D imaging stacks were morphologically segmented into regions of interest (ROIs) generally corresponding to single neurons. For each ROI, changes in GCaMP6 fluorescence intensity over time (time series) were extracted, demixed, denoised, and then z-scored. We defined an ROI as responding to a particular stimulus based on a regression coefficient and *r*^*2*^ threshold from the WT group. The coordinates of responsive ROIs from multiple animals were warped onto a reference brain (using the Z-brain atlas).

**Supplementary Figure 2.**
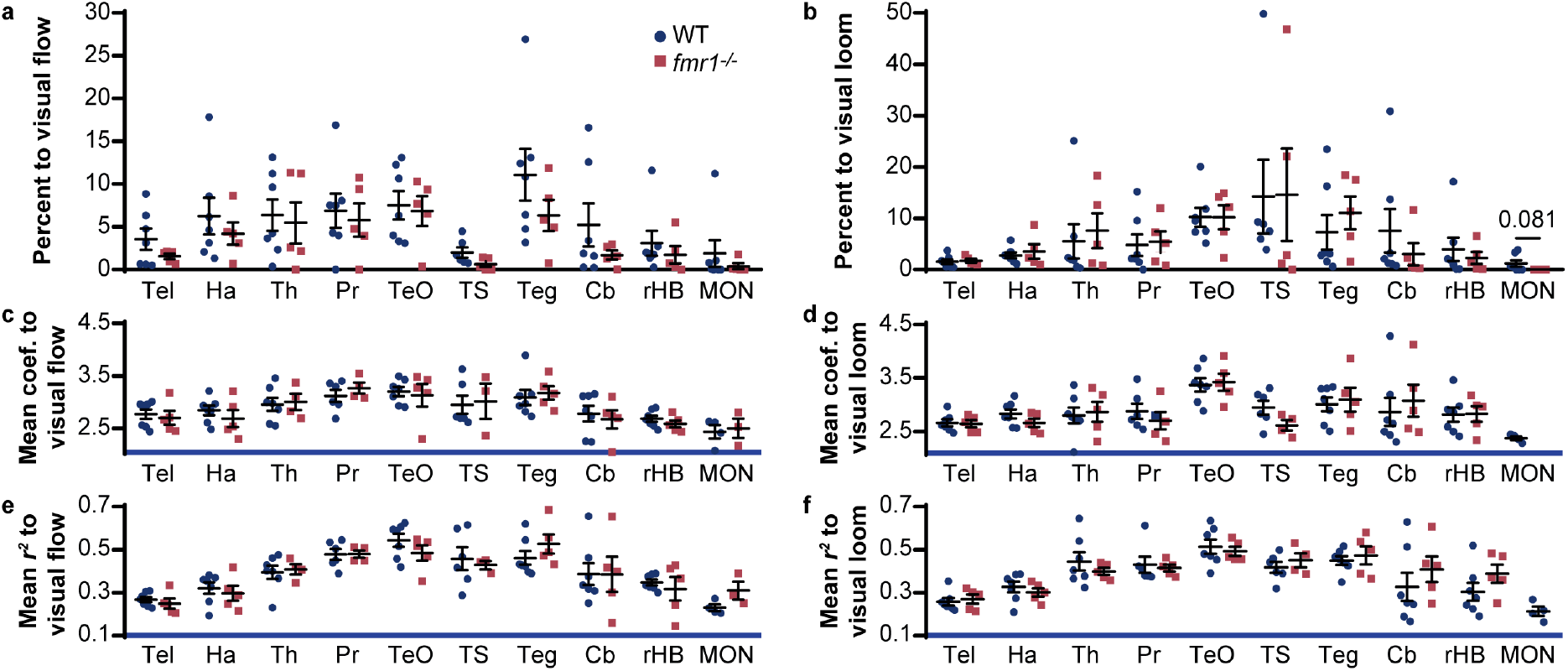
Responsiveness of brain regions to visual stimuli in WT and *fmr1*^−/−^ larvae. Percentage of ROIs in wild-type (WT) and mutant (*fmr1*^−/−^) larvae responding to visual flow **(a)** and loom **(b)** across the ten brain regions of interest (mean ± s.e.m.). Mean regression coefficient **(c, d)** and *r*^*2*^ **(e, f)** values of ROIs responding to visual flow **(c, e)** and loom **(d, f)** in these regions. Significance was tested using the Kruskal-Wallis test with Dunn’s multiple comparison test, and is only shown in cases where p < 0.1. Cb, cerebellum; Ha, habenulae; MON, medial octavolateralis nucleus; Pr, pretectum; rHB, remaining hindbrain with the Cb and MON excluded; Teg, tegmentum; TeO, optic tectum; Th, thalamus; TS, torus semicircularis.

**Supplementary Table 1.** Summary of the statistical analyses used to quantify baseline activity. The baseline dataset was analysed using the unpaired, Kruskal-Wallis test with Dunn’s multiple comparison test. Wild-type (WT) *n* = 5; heterozygous mutant (*fmr*^+/−^) *n* = 12; homozygous mutant (*fmr1*^−/−^) *n* = 7.

| *Analysis* | **WT;*fmrI*<sup>+/-</sup>**<br>**(mean ± SD)** | *P* | **WT;*fmrI*<sup>-/-</sup>**<br>**(mean ± SD)** | *P* |
| --- | --- | --- | --- | --- |
| *Mean calcium events per minute* | 3.60 ± 0.41;<br>3.40 ± 0.40 | 0.8744 | 3.60 ± 0.41;<br>3.51 ± 0.71 | >0.9999 |
| *SD of calcium events per minute* | 1.98 ± 0.19;<br>1.82 ± 0.17 | 0.1614 | 1.98 ± 0.19;<br>1.82 ± 0.12 | 0.1941 |
| *Mean correlation coefficient* | 0.12 ± 0.05;<br>0.13 ± 0.06 | >0.9999 | 0.12 ± 0.05;<br>0.13 ± 0.06 | >0.9999 |

**Supplementary Table 2.** Summary of the statistical analyses used to quantify results from the multisensory experiment. The multisensory dataset was analysed using unpaired, two-tailed Mann-Whitney tests for regions of interest (ROIs) above the responsive threshold. Wild-type (WT) *n* = 7 and homozygous mutant (*fmr1*^−/−^) *n* = 5. Cb, cerebellum; Ha, habenulae; MON, medial octavolateralis nucleus; Pr, pretectum; rHB, remaining hindbrain with the MON and Cb excluded; Teg, tegmentum; Tel, telencephalon; TeO, optic tectum; Th, thalamus; TS, torus semicircularis.

| *Analysis* | *Brain region* | *Visual flow* |  | *Visual loom* |  | *Auditory* |  |
| --- | --- | --- | --- | --- | --- | --- | --- |
|  |  | WT;*fmrI*<sup>-/-</sup><br>(mean ± SD) | *P* | WT;*fmrI*<sup>-/-</sup><br>(mean ± SD) | *P* | WT;*fmrI*<sup>-/-</sup><br>(mean ± SD) | *P* |
| *Percent responsive to stimuli* | *Whole brain* | 4.58 ± 3.09;<br>3.06 ± 1.70 | 0.6389 | 4.22 ± 3.09;<br>4.05 ± 2.62 | >0.999<br>9 | 4.41 ± 1.08;<br>8.40 ± 4.75 | 0.1490 |
|  | *Tel* | 3.55 ± 3.26;<br>1.55 ± 0.70 | 0.6389 | 1.55 ± 1.20;<br>1.72 ± 0.83 | 0.7551 | 1.23 ± 0.62;<br>1.77 ± 0.75 | 0.3434 |
|  | *Ha* | 6.26 ± 5.67;<br>4.21 ± 2.88 | 0.6389 | 2.80 ± 1.52;<br>3.54 ± 3.16 | >0.999<br>9 | 1.11 ± 0.75;<br>1.38 ± 0.74 | 0.2677 |
|  | *Th* | 6.38 ± 4.90;<br>5.46 ± 5.39 | 0.7551 | 5.52 ± 8.87;<br>7.59 ± 7.63 | 0.6389 | 18.7 ± 5.96;<br>18.7 ± 3.71 | 0.7551 |
|  | *Pr* | 6.90 ± 5.23;<br>5.78 ± 4.35 | 0.9192 | 4.76 ± 5.52;<br>5.39 ± 4.54 | 0.8763 | 9.35 ± 5.49;<br>10.4 ± 4.38 | 0.7551 |
|  | *TeO* | 7.54 ± 4.37;<br>6.85 ± 3.90 | 0.8763 | 10.2 ± 4.86;<br>10.2 ± 5.28 | 0.7551 | 1.11 ± 0.94;<br>1.89 ± 0.09 | 0.0480 |
|  | *TS* | 2.02 ± 1.48;<br>0.64 ± 0.72 | 0.1190 | 14.2 ± 17.6;<br>14.6 ± 20.1 | 0.4286 | 9.99 ± 6.22;<br>6.58 ± 4.60 | 0.5368 |
|  | *Teg* | 11.1 ± 7.99;<br>6.34 ± 4.11 | 0.3434 | 7.29 ± 8.88;<br>11.1 ± 7.20 | 0.4318 | 8.38 ± 6.54;<br>14.0 ± 7.30 | 0.2677 |
|  | *Cb* | 5.23 ± 6.59;<br>1.70 ± 1.17 | 0.4318 | 7.57 ± 11.2;<br>3.02 ± 4.87 | 0.2020 | 3.29 ± 1.59;<br>11.0 ± 9.77 | 0.1515 |
|  | *rHB* | 3.09 ± 3.82;<br>1.76 ± 2.28 | 0.5303 | 3.96 ± 6.07;<br>2.29 ± 2.66 | >0.999<br>9 | 13.8 ± 4.11;<br>24.5 ± 15.0 | 0.1061 |
|  | *MON* | 1.89 ± 4.14;<br>0.41 ± 0.78 | >0.999<br>9 | 1.21 ± 1.67;<br>0 ± 0 | 0.0808 | 18.7 ± 11.1;<br>38.6 ± 18.5 | 0.0756 |
| *Mean regression coefficient of responsive ROIs* | *Whole brain* | 2.99 ± 0.17;<br>2.98 ± 0.33 | 0.8763 | 3.18 ± 0.28;<br>3.13 ± 0.30 | 0.8763 | 3.42 ± 0.178;<br>3.30 ± 0.09 | 0.2020 |
|  | *Tel* | 2.77 ± 0.25;<br>2.70 ± 0.29 | 0.7551 | 2.66 ± 0.17;<br>2.64 ± 0.15 | >0.999<br>9 | 2.82 ± 0.10;<br>2.85 ± 0.19 | >0.999<br>9 |
|  | *Ha* | 2.84 ± 0.24;<br>2.69 ± 0.36 | 0.4318 | 2.83 ± 0.23;<br>2.66 ± 0.18 | 0.2677 | 2.97 ± 0.43;<br>2.83 ± 0.18 | 0.8763 |
|  | *Th* | 2.96 ± 0.33;<br>3.00 ± 0.31 | 0.9273 | 2.80 ± 0.39;<br>2.86 ± 0.41 | 0.8763 | 3.59 ± 0.19;<br>3.42 ± 0.13 | 0.1490 |
|  | *Pr* | 3.12 ± 0.28;<br>3.27 ± 0.21 | 0.4762 | 2.88 ± 0.35;<br>2.70 ± 0.36 | 0.5368 | 3.32 ± 0.38;<br>3.37 ± 0.26 | 0.7551 |
|  | *TeO* | 3.20 ± 0.23;<br>3.13 ± 0.48 | >0.999<br>9 | 3.37 ± 0.34;<br>3.42 ± 0.36 | 0.8763 | 3.04 ± 0.19;<br>3.03 ± 0.26 | 0.8763 |
|  | *TS* | Low *n* | - | 2.95 ± 0.32;<br>2.61 ± 0.12 | 0.1143 | 3.57 ± 0.40;<br>3.36 ± 0.21 | 0.1443 |
|  | *Teg* | 3.09 ± 0.38;<br>3.17 ± 0.29 | 0.3434 | 3.01 ± 0.34;<br>3.09 ± 0.50 | >0.999<br>9 | 3.38 ± 0.30;<br>3.26 ± 0.29 | 0.6389 |
|  | *Cb* | 2.78 ± 0.39;<br>2.67 ± 0.39 | 0.3434 | 2.89 ± 0.68;<br>3.07 ± 0.67 | 0.5303 | 3.20 ± 0.26;<br>3.07 ± 0.21 | 0.3434 |
|  | *rHB* | 2.68 ± 0.16;<br>2.59 ± 0.15 | 0.2677 | 2.81 ± 0.36;<br>2.82 ± 0.33 | 0.7551 | 3.52 ± 0.14;<br>3.40 ± 0.07 | 0.1061 |
|  | *MON* | Low *n* | - | Low *n* | - | 3.72 ± 0.34;<br>3.34 ± 0.13 | 0.0732 |
| *Mean $r^2$ of responsive ROIs* | *Whole brain* | 0.42 ± 0.03;<br>0.40 ± 0.08 | 0.8763 | 0.44 ± 0.07;<br>0.42 ± 0.04 | 0.6389 | 0.49 ± 0.04;<br>0.46 ± 0.06 | 0.2020 |
|  | *Tel* | 0.27 ± 0.03;<br>0.25 ± 0.05 | 0.4318 | 0.26 ± 0.04;<br>0.27 ± 0.05 | 0.8763 | 0.29 ± 0.04;<br>0.31 ± 0.04 | 0.6389 |
|  | *Ha* | 0.32 ± 0.07;<br>0.30 ± 0.08 | 0.5303 | 0.33 ± 0.06;<br>0.30 ± 0.04 | 0.4318 | 0.37 ± 0.08;<br>0.35 ± 0.06 | 0.8763 |
|  | *Th* | 0.39 ± 0.08;<br>0.41 ± 0.05 | >0.999<br>9 | 0.45 ± 0.11;<br>0.40 ± 0.03 | 0.6389 | 0.56 ± 0.06;<br>0.53 ± 0.04 | 0.2677 |
|  | *Pr* | 0.48 ± 0.06;<br>0.48 ± 0.03 | >0.999<br>9 | 0.43 ± 0.09;<br>0.42 ± 0.03 | 0.7922 | 0.53 ± 0.08;<br>0.51 ± 0.06 | 0.5303 |
|  | *TeO* | 0.54 ± 0.08;<br>0.48 ± 0.08 | 0.3434 | 0.51 ± 0.09;<br>0.49 ± 0.05 | 0.6389 | 0.43 ± 0.07;<br>0.41 ± 0.08 | 0.6389 |
|  | *TS* | Low *n* | - | 0.42 ± 0.06;<br>0.45 ± 0.07 | 0.6095 | 0.57 ± 0.09;<br>0.54 ± 0.01 | 0.4762 |
|  | *Teg* | 0.46 ± 0.09;<br>0.53 ± 0.10 | 0.1490 | 0.45 ± 0.05;<br>0.47 ± 0.09 | 0.5303 | 0.53 ± 0.05;<br>0.49 ± 0.09 | 0.4318 |
|  | *Cb* | 0.39 ± 0.13;<br>0.39 ± 0.18 | >0.999<br>9 | 0.33 ± 0.17;<br>0.41 ± 0.13 | 0.3434 | 0.45 ± 0.07;<br>0.42 ± 0.07 | 0.7551 |
|  | *rHB* | 0.35 ± 0.03;<br>0.32 ± 0.12 | >0.999<br>9 | 0.30 ± 0.11;<br>0.39 ± 0.09 | 0.2020 | 0.51 ± 0.05;<br>0.48 ± 0.07 | 0.6389 |
|  | *MON* | Low *n* | - | Low *n* | - | 0.59 ± 0.04;<br>0.49 ± 0.04 | 0.0025 |

**Supplementary Table 3.** Summary of the statistical analyses used to quantify auditory sensitivity. The auditory sensitivity dataset was analysed using unpaired, one-tailed Mann-Whitney tests for regions of interest (ROIs) above the responsive threshold. Wild-type (WT) *n* = 5 and homozygous mutant (*fmr1*^−/−^) *n* = 7. Cb, cerebellum; Ha, habenula; MON, medial octavolateralis nucleus; Pr, pretectum; rHB, remaining hindbrain with the MON and Cb excluded; Teg, tegmentum; Tel, telencephalon; TeO, optic tectum; Th, thalamus; TS, torus semicircularis.

| *Analysis* | *Brain region* | *-21 dB*<br>*P* | *-18 dB*<br>*P* | *-15 dB*<br>*P* | *-12 dB*<br>*P* | *-9 dB*<br>*P* | *-6 dB*<br>*P* | *-3 dB*<br>*P* | *0 dB*<br>*P* |
| --- | --- | --- | --- | --- | --- | --- | --- | --- | --- |
| *Mean regression coefficient of responsive ROIs* | *Tel* | 0.3242 | 0.4636 | 0.1576 | 0.2636 | 0.0121 | 0.0545 | 0.2061 | 0.4636 |
|  | *Ha* | 0.4217 | 0.1717 | 0.4381 | 0.1338 | 0.0745 | 0.2159 | 0.3775 | 0.3775 |
|  | *Th* | 0.0240 | 0.1338 | 0.0152 | 0.0530 | 0.0152 | 0.1717 | 0.0366 | 0.0745 |
|  | *Pr* | 0.3194 | 0.3194 | 0.4381 | 0.3775 | 0.3775 | 0.4381 | 0.0530 | 0.3775 |
|  | *TeO* | 0.4381 | 0.3194 | 0.4381 | 0.2652 | 0.0088 | 0.3775 | 0.0745 | 0.3775 |
|  | *TS* | 0.3775 | 0.3194 | 0.1717 | 0.0366 | 0.0366 | 0.2652 | 0.2159 | 0.3775 |
|  | *TS2* | 0.2061 | 0.1152 | 0.3242 | 0.4636 | 0.500 | 0.1152 | 0.1152 | 0.0364 |
|  | *Teg1* | 0.2159 | 0.2652 | 0.1717 | 0.1388 | 0.0013 | 0.3194 | 0.1717 | 0.1338 |
|  | *Cb* | 0.2652 | 0.3775 | 0.2652 | 0.1717 | 0.2159 | 0.2159 | 0.0745 | 0.3775 |
|  | *rHB* | 0.2159 | 0.2652 | 0.1338 | 0.4381 | 0.0366 | 0.3775 | 0.1010 | 0.500 |
|  | *MON* | 0.4381 | 0.4381 | 0.5000 | 0.500 | 0.3194 | 0.500 | 0.0240 | 0.2159 |

